# FapA extends the flagellar cage-dependent network into the cytoplasm to control surface-associated behaviors in *Helicobacter pylori*

**DOI:** 10.64898/2026.10.23.761471

**Authors:** Jiunn N. C. Fong, Corey D. Witt, Eric H. L. Huang, Jian Yue, Xiaolin Liu, Anna Roujeinikova, Jun Liu, Karen M. Ottemann

## Abstract

Flagellar systems enable many host-associated bacteria to colonize successfully. One such pathogen is *Helicobacter pylori*, which assembles an elaborate, high-torque flagellar motor that has a wide C-ring, 18 stator complexes, and is surrounded by a cage that includes homologs of the type IV pilus proteins PilM, PilN, and PilO. PilMNO act to repress soft-agar migration while promoting cell aggregation, microcolony, and biofilm formation, but the pathway linking the flagellar cage to these behaviors is unknown. Here we identify FapA, a member of the flagellar assembly protein A (FapA) family, as a component of this pathway. Deletion of *fapA* increased soft-agar migration and reduced microcolony and biofilm formation, similar to deletion of *pilMNO*. Epistasis analysis placed FapA downstream of the PilMNO flagellar cage. Using fluorescent fusions and protein-protein interaction experiments, FapA was found to localize to the cell poles, the sites of flagellar motor placement, and interact with the cytoplasmic PilM component of the cage. Indeed, cryo-electron tomography experiments detected cytoplasmic density near PilM that was lost in the Δ*fapA* deletion mutant. Phylogenetic analysis suggests that FapA proteins are more widespread than previously appreciated, being found in multiple bacterial phyla including Campylobacterota, Bacillota, Spirochaetota, Desulfobacterota, Thermotogota, Synergistota, and Pseudomonadota. Together, these findings support that the *H. pylori* flagellar cage has additional components that reside in the cytoplasm but similarly function to limit motility and activate a sessile/biofilm lifestyle. Given the widespread nature of FapA, our results suggest that many other bacteria might employ FapA homologs for similar purposes.

**Importance:** The flagella of *Helicobacter pylori* enable it to colonize the stomach, but its flagellar motor appears to do more than propel the bacterium. A protein cage surrounding the motor helps determine whether cells swim or settle into microcolonies and biofilms, yet the molecular components that link this structure to cellular behaviors remain poorly understood. This knowledge gap is particularly intriguing because *H. pylori* lacks the cyclic-di-GMP signaling system that many bacteria use to coordinate motility and biofilm development. Our identification of FapA as a cytoplasmic component of the cage-dependent regulatory network provides a foundation for understanding how *H. pylori* coordinates motility and surface-associated growth and given the widespread nature of FapA, raises the possibility that many other bacteria use similar mechanisms.

## Introduction

Flagellar motility enables many host-associated bacteria to colonize specific host niches. The bacterial flagellum is a nanoscale rotary motor composed of a filament, hook, and basal body that converts ion-motive force into torque to propel the cell through its environment (Colin et al. 2021). Motility is particularly important for *Helicobacter pylori*, a gastric pathogen that infects approximately half of the global population (Crowe 2019; Shah et al. 2023; Malfertheiner et al. 2023). To establish and maintain infection, *H. pylori* must escape the acidic lumen and penetrate the thick gastric mucus layer to reach the gastric glands, where the pH is closer to neutral (Celli et al. 2009; Keilberg et al. 2016; Schreiber et al. 2004). Colonization therefore depends on flagellar motility: *H. pylori* mutants with impaired motility are markedly deficient in establishing infection in diverse animal models (Eaton et al. 1996; Foynes et al. 1999; Ottemann and Lowenthal 2002).

*H. pylori* assembles structurally elaborate flagellar motors containing numerous accessory proteins absent from the canonical motors of *Escherichia coli* and *Salmonella enterica* (Qin et al. 2017; Liu, Tachiyama, et al. 2024; Shoichi Tachiyama et al. 2025; Gao et al. 2014; Zhou and Roujeinikova 2021; Fong et al. 2026). Its motor incorporates 18 stator units, among the highest numbers reported in bacteria, together with a widened C-ring and additional periplasmic scaffolds (Chaban et al. 2018; Qin et al. 2017; Liu, Tachiyama, et al. 2024; Shoichi Tachiyama et al. 2025; Zhou and Roujeinikova 2021). These features are hallmarks of the high-torque flagellar motors of the Campylobacterota (Feng et al. 2026; Chaban et al. 2018; Mo et al. 2022), an architecture well suited for movement through viscous gastrointestinal mucus (Beeby et al. 2016; Lertsethtakarn et al. 2011; Burnham and Hendrixson 2018).

Among these accessory structures is a unique flagellar cage that surrounds the motor, first identified by cryo-electron tomography (cryo-ET) (Qin et al. 2017). Subsequent work determined the inner-membrane-proximal part of the cage to be composed of PilM, PilN, and PilO, remote homologs of proteins involved in T4P biogenesis (Liu, Tachiyama, et al. 2024; Sagoo et al. 2024; Bonny et al. 2024). Mutants lacking cage components retain flagella and migrate farther through soft agar, despite showing no increase in swimming speed relative to wild type (Liu, Tachiyama, et al. 2024; Sagoo et al. 2024; Bonny et al. 2024). They also form smaller microcolonies and exhibit delayed biofilm initiation, suggesting that the cage balances motility with surface attachment (Liu, Tachiyama, et al. 2024). These phenotypes suggest that the cage influences cellular behavior beyond its role affecting swimming at the motor. Counterclockwise flagellar rotation promotes biofilm initiation only when the cage is intact, placing the cage downstream of rotation as a potential signal-transducing module (Liu, Lertsethtakarn, et al. 2024). The molecular pathway underlying these behaviors, however, remains unclear. Many bacteria coordinate motility and biofilm formation through the second messenger cyclic-di-GMP (c-di-GMP), but *H. pylori* has no known c-di-GMP signaling systems and no c-di-GMP was detected (Galperin 2018; Liu, Tachiyama, et al. 2024; Orr et al. 2016). Identifying proteins associated with the cage may therefore reveal an undiscovered c-di-GMP-independent mechanism that balances motile and sessile lifestyles in *H. pylori*.

Here, we identified HPG27_837 (HP0884 in 26695 strain), a hypothetical protein previously co-immunoprecipitated with the stator protein MotB (Liu, Tachiyama, et al. 2024), as a component of the pathway linking the flagellar cage to soft-agar migration and biofilm formation. The same co-immunoprecipitation experiment also recovered PilM, PilN, and PilO, suggesting a potential association between HPG27_837 and the flagellar cage. HPG27_837 is annotated in UniProt as a DUF342 domain-containing protein with similar predicted structure as *Vibrio vulnificus* flagellar assembly protein A (FapA) and hereafter referred to as FapA. *Vibrio vulnificus* FapA has been suggested to contribute to polar flagellar assembly (Park et al. 2016, 2019) and contains a β-solenoid domain as annotated in UniProt. The two proteins share approximately 14% amino acid sequence identity in a ClustalW pairwise alignment (Larkin et al. 2007). We show that, unlike in *V. vulnificus*, deletion of *fapA* in *H. pylori* does not result in aflagellation but instead increases soft-agar migration and reduces microcolony and biofilm formation, phenotypes that resemble those observed upon deletion of *pilMNO*. Epistasis analyses further suggest that FapA acts downstream of PilMNO in the pathway regulating these behaviors. We also show that FapA localizes to the cell poles, the location of the flagellar motors, and interacts directly with PilM, the cytoplasmic cage component. Cryo-ET and subtomogram averaging revealed a density near PilM and the flagellar stators, likely corresponding to FapA. Together, these findings suggest that FapA may act as a downstream effector of the flagellar cage, extending its influence into the cytoplasm to modulate motility and cellular behavior in *H. pylori*.

## Results

### FapA and the flagellar cage promote biofilm formation and restrict soft-agar migration in *H. pylori*

Previous work had shown that co-immunoprecipitation with the flagellar stator protein MotB identified FapA (HPG27_837) as a highly enriched protein (Liu, Tachiyama, et al. 2024). To investigate its function, we generated multiple independent Δ*fapA* mutant strains alone and in combination with cage mutants, *ΔpilM* and *ΔpilMNO,* and assessed biofilm formation and soft-agar migration of the resulting strains (Fig. 1, Fig. S1). We previously showed that Δ*pilM* and Δ*pilMNO* migrate faster in soft agar, also called semi-solid agar, and form less biofilm than wild type (Liu, Tachiyama, et al. 2024). The Δ*fapA* strain phenocopied Δ*pilM* and Δ*pilMNO* strains, exhibiting enhanced migration through soft agar (Fig. 1A and B) and reduced biofilm formation after 2 days (Fig. 1C) relative to the wild-type G27 parent. Adding Δ*fapA* to *ΔpilM* or Δ*pilMNO* mutants did not produce a statistically detectable further change in migration or biofilm biomass, suggesting that FapA acts downstream of the cage pathway (Fig. 1A and C). However, the less severe phenotypes of the Δ*fapA* strain compared to those of Δ*pilMNO* suggest that FapA accounts for only part of the cage-dependent output, implying the possible involvement of additional pathways. Complementation of the Δ*fapA* strain migration phenotype by addition of chromosomal copy of *fapA*-mNeonGreen, placed in an intergenic region, restored Δ*fapA* soft-agar migration to wild-type levels (Fig. S1).

**Fig. 1.**
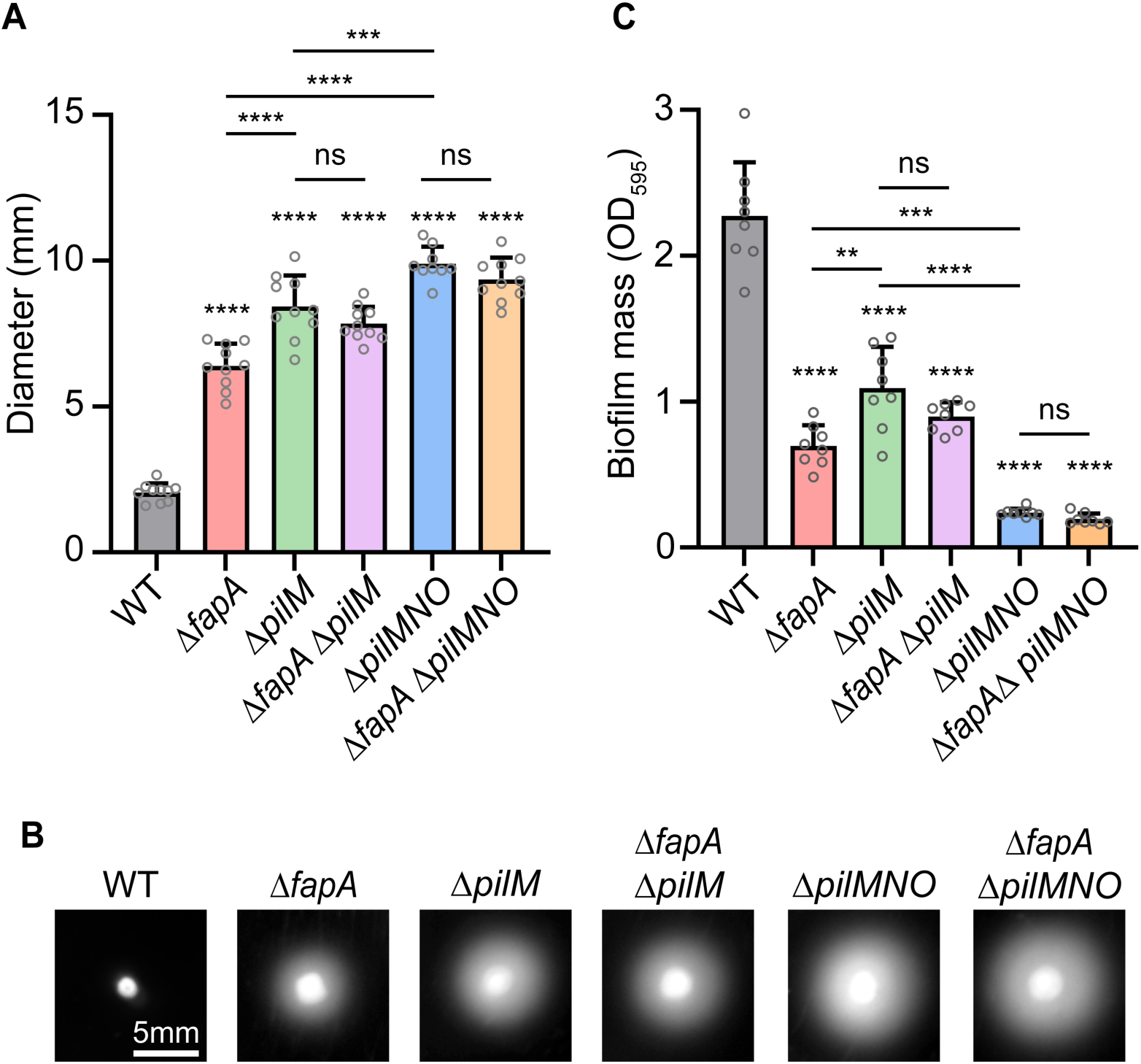
FapA phenocopies PilMNO in repressing soft-agar motility and promoting biofilm formation. (A) Quantification of soft-agar migration, (B) representative images of migration zones, and (C) quantification of biofilm biomass. Compared with G27 wild type (WT), Δ*fapA* and the cage mutants exhibited increased migration zone diameters and decreased biofilm formation after two days of growth. Each data point represents a technical replicate from a single biological replicate. Experiments were performed with at least three independent biological replicates. Statistical significance was determined using one-way ANOVA followed by Tukey’s multiple-comparisons test. Error bars represent standard deviations. ****, *p* <0.0001; ***, *p* = 0.0007; **, *p* = 0.004; ns, not significant.

We previously found that PilO restricts soft-agar migration independently of flagellar rotational bias (Liu, Tachiyama, et al. 2024). To test whether FapA also functions independently in these backgrounds, we introduced Δ*fapA* into strains with clockwise bias (Δ*cheZ)* or counterclockwise bias (Δ*cheV1*). Significant differences were observed between the corresponding single and double mutants (Δ*cheV1* vs Δ*fapA* Δ*cheV1*, and Δ*cheZ* vs Δ*fapA* Δ*cheZ*), indicating that, like PilO, FapA normally represses soft-agar migration independently of flagellar rotational bias (Fig. S1).

We observed that biofilms in *H. pylori* G27 cultures frequently form as pellicles (floating biofilms). The crystal violet assay captures only surface-associated biomass, which may introduce variability and reduce sensitivity for strains with subtle differences in biofilms. We therefore examined cells grown on soft-agar pads, allowing a higher-resolution assessment of biofilm formation in terms of cell aggregation and microcolony development, while simultaneously observing migration behavior (Fig. 2). After one day of growth, the wild-type strain exhibited the highest microcolony density, with the Δ*fapA*, Δ*pilM*, and Δ*pilMNO* strains exhibiting progressively lower microcolony formation that were apparent by visual examination (Fig. 2A, middle row) and quantitative image analysis (Fig. 2B). The Δ*fapA* Δ*pilM* and Δ*fapA* Δ*pilMNO* double mutants did not differ significantly from their parental single mutants Δ*pilM* and Δ*pilMNO* (Fig. 2B), further supporting the idea that FapA functions downstream of PilMNO. Both the Δ*fapA* and Δ*pilM* strains exhibited less microcolony formation compared to wild type (Fig. 2B), in a manner that varied in terms of which was lower compared to the crystal violet assay (Fig. 1C), consistent with the idea that these assays capture different aspects of aggregative behavior. In sum, this assay allowed us to order the strains based on microcolony density, from highest to lowest: G27 > Δ*fapA* > Δ*pilM* and Δ*fapA* Δ*pilM* > Δ*pilMNO* and Δ*fapA* Δ*pilMNO*.

**Fig. 2.**
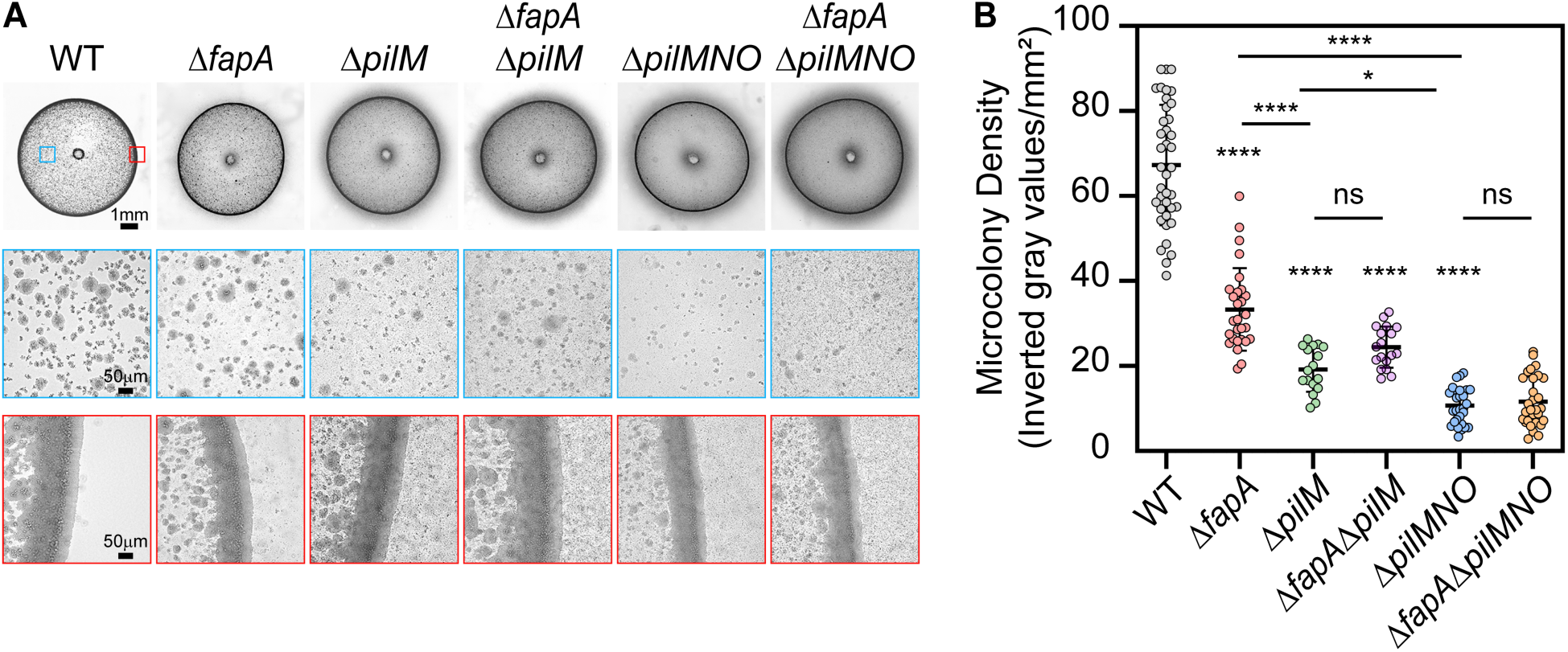
FapA and PilMNO promote cell aggregation and microcolony formation. (A) Representative images of microcolonies and (B) quantification of microcolony density of G27 wild type (WT), Δ*fapA*, Δ*pilM*, Δ*fapA* Δ*pilM*, Δ*pilMNO* and Δ*fapA* Δ*pilMNO* after one day of growth on soft-agar pads, illustrating the positive roles of FapA and PilMNO in cell aggregation and microcolony formation. In A, the top row shows the entire culture spot, while the middle and bottom rows show microcolonies within the spot and cells migrating beyond the spot edge, respectively. Blue and red boxes (in A) indicate the corresponding regions shown at higher magnification in the middle and bottom rows, respectively. Each data point (in B) represents a technical replicate from at least three biological replicates. Statistical significance was determined using one-way ANOVA followed by Tukey’s multiple-comparisons test. Error bars represent standard deviations. ****, *p* <0.0001; *, *p* = 0.0271; ns, not significant.

Migration behavior in the soft-agar pads followed a trend that was inverse to the microcolony density, with all mutants showing dispersal beyond the spot edge compared to wild type (Fig. 2A, bottom row). Few single cells were observed beyond the spot edge of G27 wild type, consistent with low migration in wild-type cells. In contrast, mutant strains exhibited higher single-cell and microcolony density beyond the spot edge (Fig. 2A, bottom row), in agreement with the soft-agar migration assay (Fig. 1A and B).

To explore the biofilm phenotypes further, scanning electron microscopy (SEM) was performed. This analysis further showed that Δ*fapA* strains exhibited a biofilm phenotype intermediate between wild-type G27 and the Δ*pilMNO* cage mutant. These biofilms, which were grown on glass coverslips, showed elaborate three-dimensional structures in wild-type G27, smaller microcolonies in the Δ*fapA* strain, and mostly single cells and small clusters in the *ΔpilMNO* mutant (Fig. S2), results that were overall consistent with crystal violet (Fig. 1C) and microcolony soft-agar pad assays (Fig. 2). SEM imaging further confirmed that the Δ*fapA* and Δ*pilMNO* mutant strains were flagellated, as expected, suggesting that their attenuated biofilm phenotypes result not from loss of flagellation but from disruption of a flagellar cage/FapA-dependent pathway.

Together, these data indicate that Δ*fapA* mutants phenocopy cage mutants lacking PilMNO. These mutants exhibit decreased biofilm formation and increased migration, consistent with the idea that the proteins normally promote biofilm and microcolony formation and repress soft-agar migration. In addition, the data suggest FapA functions downstream of the cage pathway and likely forms part of the flagellar cage network.

### FapA interacts with the cage protein PilM

Because double-mutant phenotypes are consistent with FapA functioning downstream of PilMNO, we tested whether FapA physically associates with proteins of the PilMNO cage. FapA is predicted to be cytoplasmic, making the cytoplasmic cage protein PilM a candidate interaction partner. Using the Bacterial Adenylate Cyclase Two-Hybrid (BACTH) system and making all combinations for FapA and PilM fusions, we found that six of eight FapA and PilM fusion combinations showed significantly elevated β-galactosidase activity relative to the negative control (Fig. 3). These findings further strengthen the notion that FapA is part of the flagellar cage network via interactions with PilM, potentially as a downstream cytoplasmic effector of cage-dependent regulation.

**Fig. 3.**
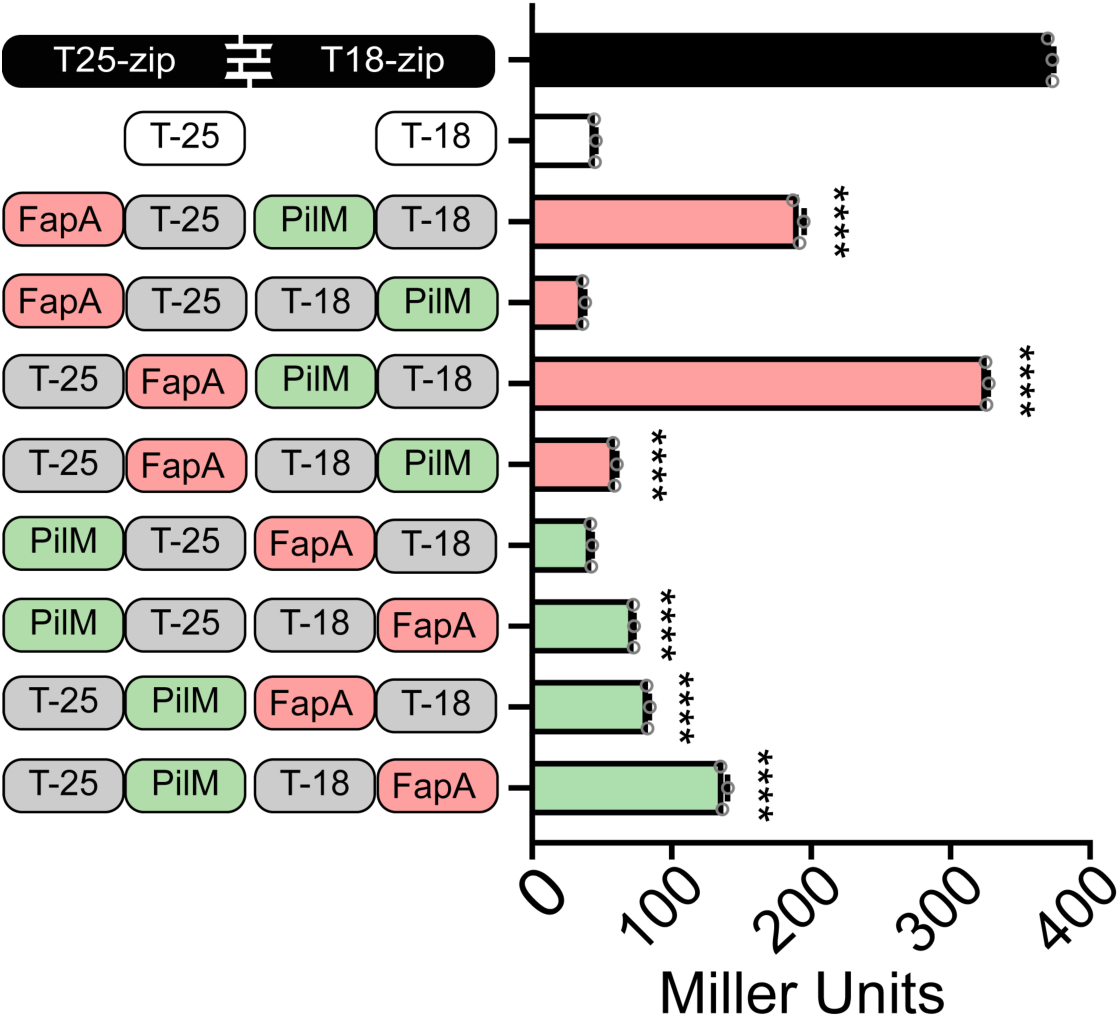
BACTH assays reveal direct interactions between FapA and the cage protein PilM. Schematics of various BACTH plasmid combinations are shown on the left, including positive (black) and negative (white) controls. BACTH assays demonstrate direct interactions between FapA and PilM, with six constructs showing significantly elevated signals compared with the negative control. β-galactosidase activity, reported as Miller Units, is shown on the right. Each data point represents a technical replicate from a single biological replicate. The assay was repeated with four biological replicates. Statistical significance was determined using one-way ANOVA followed by Tukey’s multiple-comparison test. Error bars represent standard deviations. ****, *p* <0.0001.

Given the proximity of the flagellar stator and the rotor C-ring to the flagellar cage, we used the BACTH assay in *E. coli* to test whether FapA interacts with the stator protein MotA or the rotor protein FliG. We detected no significant β-galactosidase activity when using FapA with either full-length MotA, the cytoplasmic fragment of MotA (V82-G111), or full-length FliG under the conditions tested (data not shown). This result does not exclude the possibility of these interactions in *H. pylori*, because the fusion constructs tested in *E. coli* may have steric constraints that preclude binding, or the interactions may be dynamic or transient and fall below the assay’s detection limit.

### FapA localizes to the flagellar poles

The direct interaction between FapA with the cage protein PilM suggested that FapA might localize to the flagellar motors. To test this possibility, we generated strains expressing mNeonGreen-PilO (mNG-PilO) or FapA-mNeonGreen (FapA-mNG) N- and C-terminal translational fusions, respectively, and examined their subcellular distributions. As expected, mNG-PilO localized to the cell poles in the wild-type G27 (Fig. 4A), consistent with previous observations with GFP-PilO (Liu, Tachiyama, et al. 2024). FapA-mNG localized to the cell poles in both wild-type and Δ*fapA* cells (Fig. 4A and S3), a pattern similar to that of mNG-PilO, whereas unfused NeonGreen (mNG) diffusely distributed throughout the cell (Fig. 4A). FapA-mNG retained its polar localization in clockwise-biased Δ*cheZ* and counterclockwise-biased Δ*cheA* strains, suggesting that changes in flagellar rotation bias do not disrupt FapA polar localization (Fig. S3). The FapA-mNG fusions retained function, as Δ*fapA* strains expressing these FapA-mNG fusions exhibited soft-agar migration comparable with wild-type G27 (Fig. S1).

**Fig. 4.**
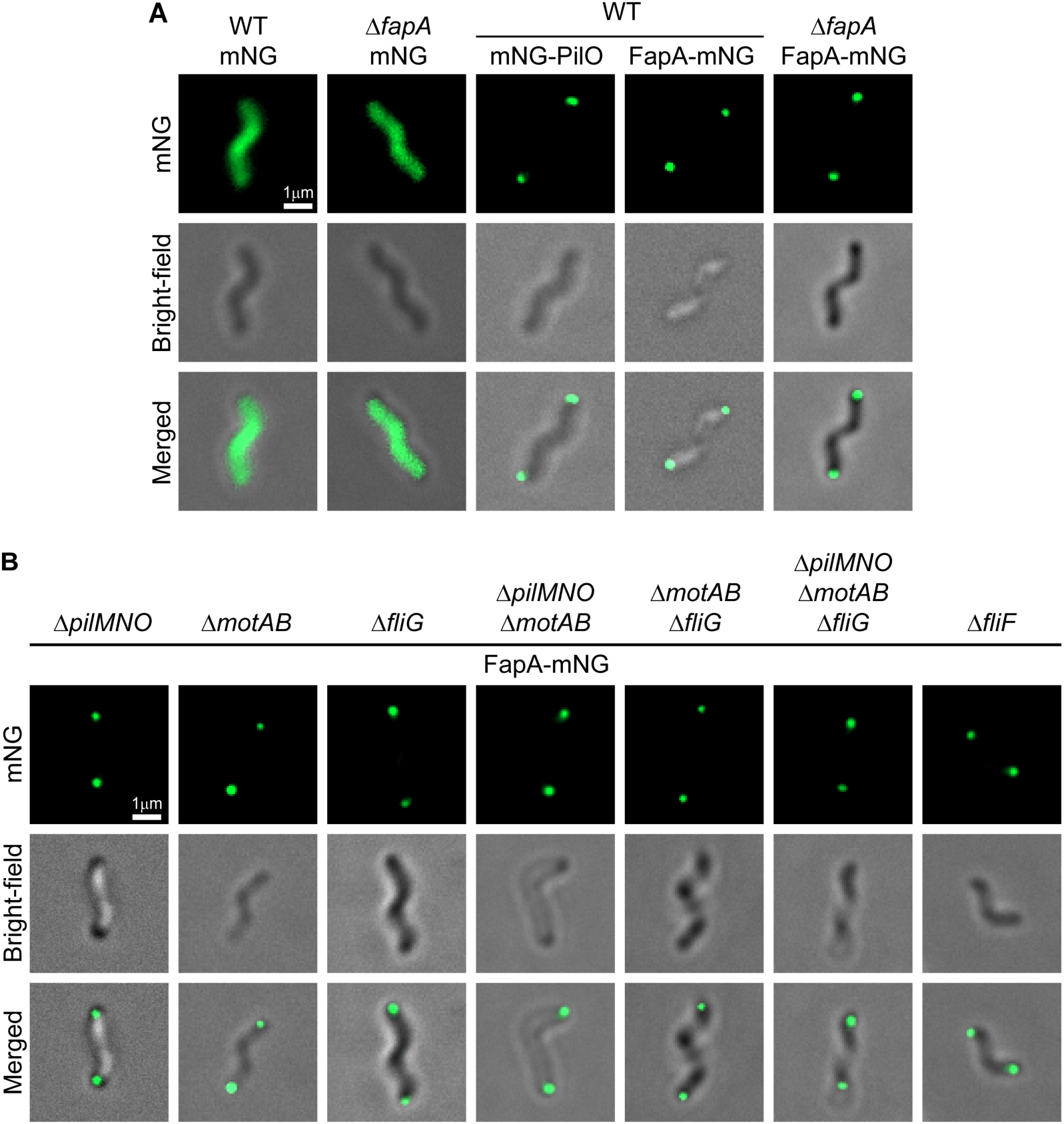
FapA localizes to the cell poles. (A) G27 wild-type (WT) strains expressing mNeonGreen-PilO (mNG-PilO) or FapA-mNeonGreen (FapA-mNG) translational fusions exhibit polar localization. FapA-mNG also retains its polar localization in the Δ*fapA* background. In contrast, the negative control, mNeonGreen (mNG), is diffusely distributed throughout the cells without forming puncta in G27 wild-type and Δ*fapA* backgrounds. (B) FapA localization in strains lacking the PilMNO cage (Δ*pilMNO*), stator (Δ*motAB*), rotor protein FliG (Δ*fliG*), or MS-ring (Δ*fliF*), and in strains carrying combined deletions of stator, motor, and/or flagellar cage components (Δ*pilMNO* Δ*motAB*, Δ*motAB* Δ*fliG* and Δ*pilMNO* Δ*motAB* Δ*fliG*). Images shown are representative of at least three biological replicates.

Because FapA interacted with PilM in the BACTH assay, we hypothesized that loss of PilM would disrupt FapA’s polar localization. Surprisingly, FapA-mNG retained polar localization in the Δ*pilMNO* mutant, a strain that lacks all three inner-membrane-proximal cage proteins (Fig. 4B). PilMNO are therefore not required for FapA polar localization, and we hypothesized there may be other factors that target FapA to the poles. The observed two-hybrid interaction of FapA with PilM suggests it may be localized near the cytoplasmic part of the cage, which is in turn near the rotor protein FliG and the stator proteins MotAB. We therefore next examined FapA-mNG localization in strains lacking Δ*fliG* or *ΔmotAB*, alone or in combination with the PilMNO cage. FapA-mNG retained polar localization in strains lacking the stator (Δ*motAB*), the rotor protein FliG (Δ*fliG*), as well as in strains lacking these components in combination with the flagellar cage (Δ*pilMNO* Δ*motAB*, Δ*motAB* Δ*fliG,* and Δ*pilMNO* Δ*motAB* Δ*fliG)* (Fig 4B). In addition, FapA-mNG remained polarly localized in a Δ*fliF* mutant lacking the flagellar MS-ring. FliF forms the base of the flagellum, is the first step of flagellar biosynthesis, and FliF mutants do not form flagella (Kubori et al. 1992; Allan et al. 2000). Together, these results indicate that FapA polar localization persists without the PilMNO cage, stator, rotor, and MS-ring. The persistence of FapA polar localization in these mutants suggests that other factors, which have yet to be identified, may help localize FapA at the cell poles.

### Deletion of *fapA* alters stator-associated densities while preserving the overall flagellar motor architecture

To provide more insight into how FapA interacts with the *H. pylori* flagella, we used cryo-ET and subtomogram averaging to compare flagellar motors in wild-type and Δ*fapA* cells. Deletion of *fapA* did not appear to alter flagellar motor number, and the mutant motors retained an overall architecture similar to that of wild-type motors (Fig. 5A and D). In particular, the disk, C-ring, and cage remained discernible, suggesting that FapA is not essential for assembly of these major structural features. Cross-sections through the cage further showed that its circumferential organization was largely preserved in the Δ*fapA* mutant (Fig. 5B and E). In contrast, densities in the stator regions were less well resolved in the mutant compared with those in wild-type motors (Fig. 5C and F). This observation suggests that FapA may influence stator assembly or occupancy within the motor. This phenotype shares features with the structural alterations previously reported for PilM mutants, which indicate that PilM contributes to maintaining the cytoplasmic architecture of the cage and stator organization (Liu, Tachiyama, et al. 2024). These similarities are consistent with the detected FapA–PilM interaction.

**Fig. 5.**
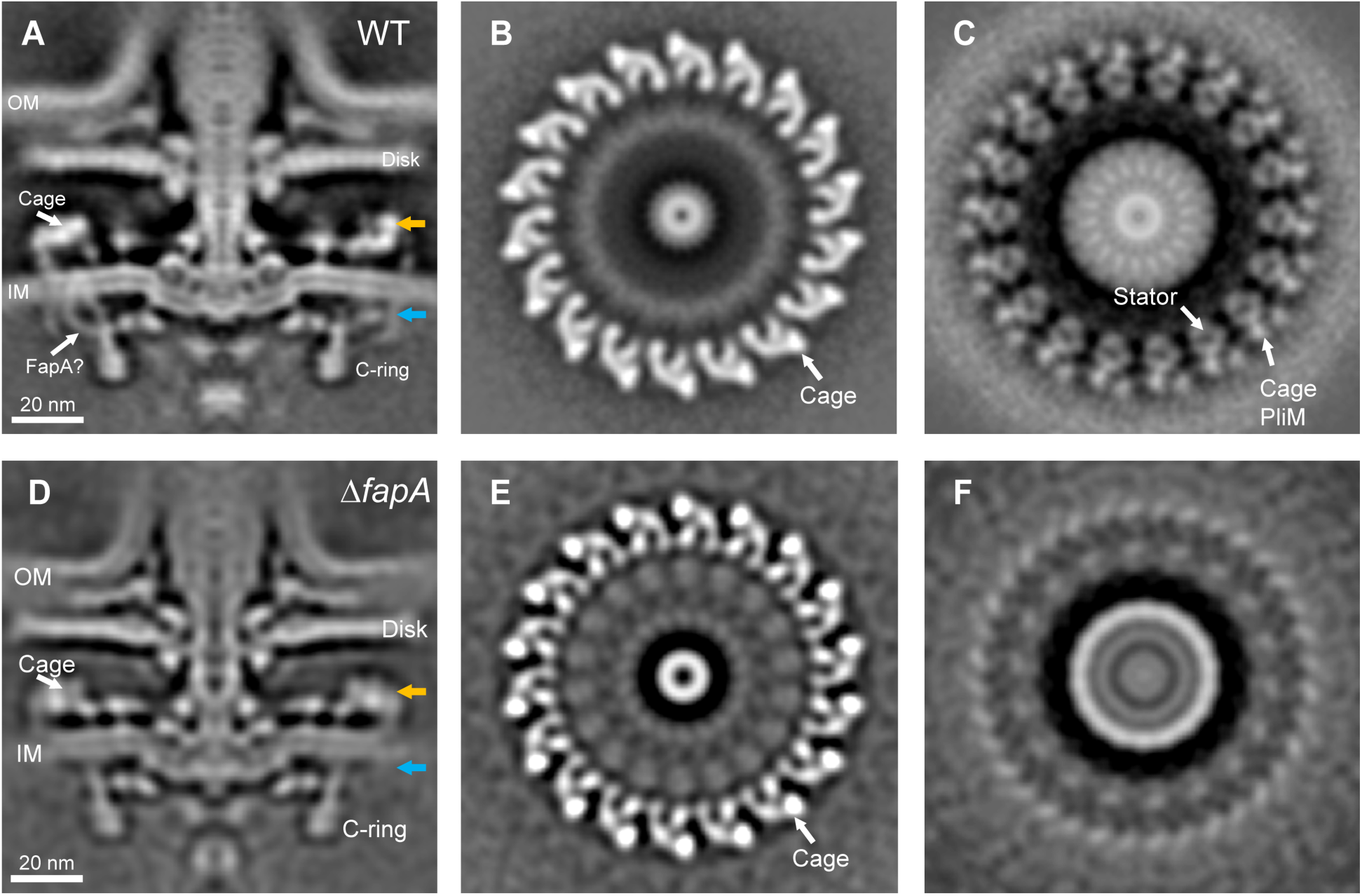
Cryo-ET analysis of *H. pylori* flagellar motors in wild-type and Δ*fapA* cells. (A) A medial slice of the wild-type (WT) motor structure shows the disk, cage, and C-ring. A putative FapA-associated density is indicated in the cytoplasmic region beneath the stator MotA. (B) A cross-section of the WT motor at the level indicated by the yellow arrow in A shows the cage surrounding the central motor axis. (C) A cross-section at the level indicated by the blue arrow in A shows the stator and peripheral cage-associated densities. The PilM density adjacent to the stator is indicated. (D) A medial slice of the motor in Δ*fapA* cells. (E, F) Corresponding cross-sections of the Δ*fapA* motor at the levels indicated by the yellow and blue arrows in D, respectively. The periplasmic portion of the cage remains visible, whereas densities in the stator region are less well resolved than in the WT motor, which may reflect greater conformational variability in this region. OM, outer membrane; IM, inner membrane. Scale bar, 20 nm.

In the wild-type reconstruction, an additional density was visible beneath the cytoplasmic portion of the stator, near the PilM-associated region (Fig. 5A). This density may correspond to FapA, although its molecular identity remains to be established. Together, these observations suggest that FapA contributes to the structural organization of the cytoplasmic stator-associated region but is dispensable for assembly of the overall motor architecture.

### *H. pylori* FapA differs from FapA homologs in other flagellated bacterial species

FapA has been reported in two other bacteria, *V. vulnificus* and *B. burgdorferi* (Zamba-Campero et al. 2024; Park et al. 2016), where it contains a C-terminal β-solenoid domain annotated as PF03961 in UniProt. These proteins also contain a FapA N-terminal region annotated as PF20250 (Fig. S4A). In contrast, PF03961 is not annotated in the *H. pylori* and *C. jejuni* FapA homologs. However, their predicted structures have a similar fold with extensive β-strand-rich regions resembling the β-solenoid regions of *V. vulnificus* and *B. burgdorferi* FapA (Fig. S4B). To examine the distribution of this domain, we searched proteomes from a database of representative flagellated bacteria (Feng et al. 2026) using the PF03961 profile hidden Markov model (HMM) from the Pfam database. We identified numerous proteins containing the domain across multiple phyla, including Pseudomonadota and Spirochaetota, the phyla to which *V. vulnificus* and *B. burgdorferi* belong, respectively (Fig. 6A). *V. vulnificus* was not in the database, but its relatives *V. cholerae* and *V. alginolyticus* were identified as containing proteins with the PF03961 β-solenoid domain. In contrast, the PF03961 profile detected relatively few proteins in Campylobacterota (Fig. 6B).

**Fig. 6.**
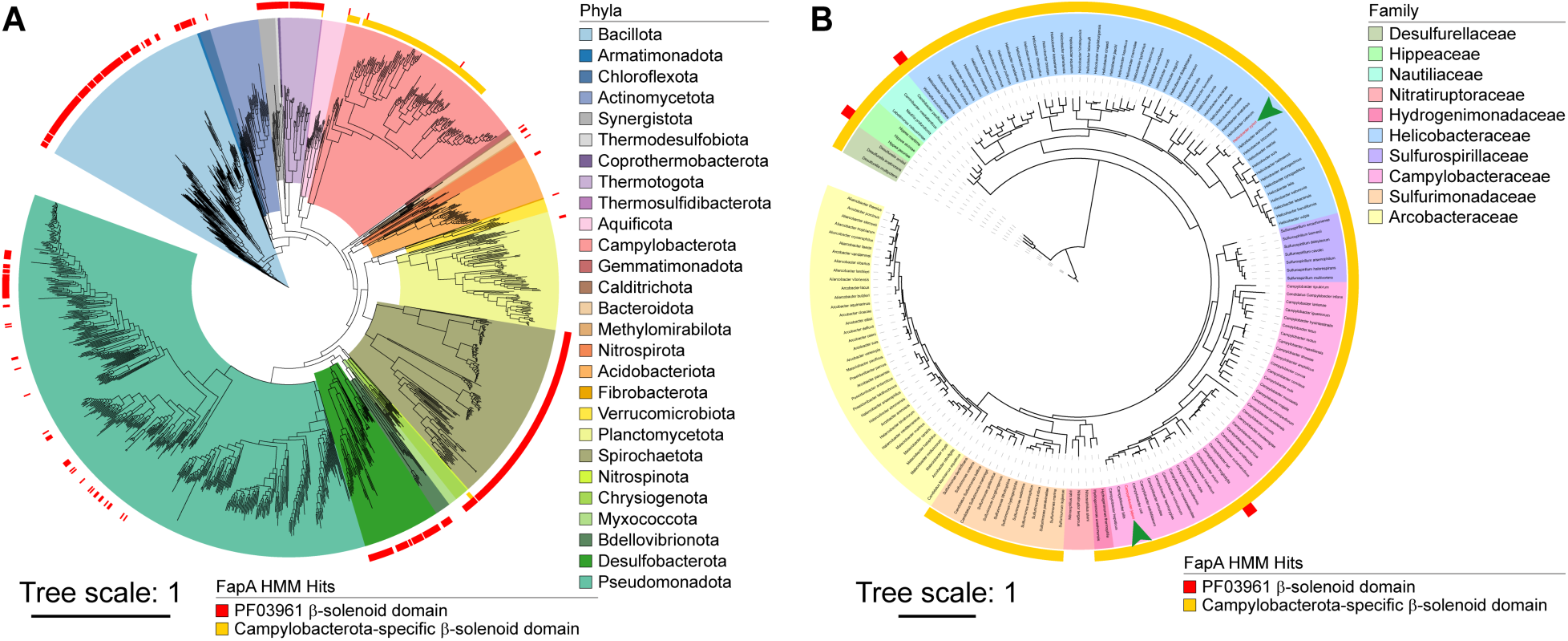
Phylogenetic distribution of FapA homologs across the represented flagellated bacteria. Distribution of FapA homologs across (A) the represented flagellated bacteria and (B) the Campylobacterota phylum. While PF03961 β-solenoid domain-containing proteins representing the FapA family proteins are widely distributed in many flagellated bacteria, Campylobacterota-specific putative β-solenoid domain-containing proteins are only found in Campylobacterota. Species names for *H. pylori* and *C. jejuni* are highlighted in red and marked with green arrowheads in B.

The annotated PF03961 regions of *V. vulnificus* and *B. burgdorferi* FapA are predicted to comprise a β-solenoid domain, an α-helical hairpin subdomain and a small β-rich subdomain at the C-terminus (Fig. S4B). We therefore used residues 300–611 of *H. pylori* FapA, which span the corresponding predicted regions, as the query to search for homologous sequences and construct a custom HMM (Fig. S4B). The HMM identified numerous additional candidate FapA homologs, predominantly in Campylobacterota, including those from *H. pylori* (*Helicobacteraceae)* and *C. jejuni* (*Campylobacteraceae*) (Fig. 6B). These findings suggest that Campylobacterota FapA proteins form a divergent subgroup within the broader FapA family, one whose putative C-terminal β-solenoid-containing region the canonical PF03961 profile fails to detect.

In *V. vulnificus*, glucose has been reported to suppress flagellar biosynthesis by promoting the sequestration of FapA from the flagellar pole by EIIA^Glc^ of the phosphoenolpyruvate:sugar phosphotransferase (PTS) system, thereby inhibiting soft-agar migration (Park et al. 2016). Given the possibility that *H. pylori* FapA represents a divergent subgroup within the FapA protein family, we hypothesized that FapA regulates soft-agar migration in *H. pylori* independently of glucose availability, unlike the glucose-dependent regulation reported in *V. vulnificus*. Indeed, wild-type *H. pylori* exhibited similar migration through soft agar in the presence and absence of supplemental glucose on days 2 and 3 (Fig. S4C). Moreover, the enhanced migration of the Δ*fapA* strain relative to the wild-type parent persisted under both conditions. These findings suggest that glucose does not substantially affect soft-agar migration in *H. pylori* under the conditions tested and that FapA-mediated repression of migration is independent of glucose availability. Together, these observations raise the possibility that FapA regulates migration through distinct mechanisms in *H. pylori* and *V. vulnificus*, potentially reflecting the divergence of *H. pylori* FapA from other members of the FapA protein family.

## Discussion

Flagella are widespread in bacteria, with conserved flagellar genes present in approximately 40% of >30,000 genomes (Andrianova et al. 2025), and they contribute to colonization and, therefore, virulence in many pathogens. Bacteria with high-torque flagellar motors such as *H. pylori* and *C. jejuni* have elaborate motor architectures and accessory proteins, including a flagellar cage, that are absent from the canonical motors of *E. coli* and *S. enterica* (Beeby et al. 2016; Shoichi Tachiyama et al. 2025; Liu, Tachiyama, et al. 2024; Feng et al. 2026; Fong et al. 2026). The flagellar cage is composed of the T4P-homologous proteins PilM, PilN and PilO, and appears to be restricted to flagellated Campylobacterota (Feng et al. 2026). It is located at the outermost edge of the basal body, surrounding the stators and extending from the cytoplasm to the cell wall (Liu, Tachiyama, et al. 2024; Fong et al. 2026; Sagoo et al. 2024; Qin et al. 2017). Δ*pilMNO* mutants exhibit increased soft-agar migration and attenuated initial biofilm formation (Liu, Tachiyama, et al. 2024), suggesting that the cage contributes to a transition between motile and sessile lifestyles. Our findings place FapA as a downstream cytoplasmic effector of the flagellar cage that extends the cage’s regulatory influence into the cytoplasm to control surface-associated behaviors.

The results presented here suggest that FapA is part of the flagellar cage network and acts downstream of the cage, phenocopying the cage mutant in inversely regulating soft-agar migration and biofilm formation/cell aggregation (Fig. 1 and 2). Consistent with this model, Δ*fapA* strains, like those lacking Δ*pilMNO*, exhibit enhanced soft-agar migration but reduced biofilm formation and cell aggregation. However, the Δ*fapA* phenotypes are less pronounced than those of Δ*pilMNO*, and combining the mutations did not produce a stronger phenotype than the Δ*pilMNO* mutant alone. One possibility to explain these findings is that the cage may also act through FapA-independent pathways. One option is a parallel pathway through PilO. In *Pseudomonas aeruginosa*, the T4P protein PilO interacts with the diguanylate cyclase SadC to regulate intracellular c-di-GMP levels, thereby modulating motility and biofilm formation (Webster et al. 2021). However, *H. pylori* genomes do not appear to encode proteins involved in c-di-GMP metabolism (Galperin 2018; Orr et al. 2016), and c-di-GMP was not detected in *H. pylori* (Liu, Tachiyama, et al. 2024). Thus, PilO may instead interact with other effector proteins to regulate motility and biofilm formation independently of FapA.

FapA localizes to the cell poles, the site of flagellar motor assembly, and directly interacts with PilM, the cytoplasmic flagellar cage component (Fig. 3 and 4). No interactions were detected between FapA and MotA or FliG in BACTH assays under the conditions tested. However, dynamic or transient contacts, as well as steric constraints in these fusion constructs, cannot be ruled out. In *V. vulnificus*, FapA polar localization is dependent on the polar landmark protein HubP (Park et al. 2019). In contrast, although *H. pylori* FapA interacts with the cage protein PilM, its polar localization is independent of the PilMNO cage, stator, rotor, and MS-ring. The persistence of polar FapA localization in Δ*fliF* and Δ*pilMNO* Δ*motAB* Δ*fliG* strains suggests that other factors contribute to FapA localization. No HubP homolog has been identified in *H. pylori* (data not shown) and whether a HubP/FimV-like protein contributes to FapA localization in *H. pylori* remains to be investigated. The molecular mechanism by which FapA modulates soft-agar migration and biofilm formation also remains unresolved. Identifying additional interacting partners, for example by co-immunoprecipitation and mass spectrometry, may help identify how FapA is recruited to the flagellar motor and how it functions in *H. pylori*.

Cryo-ET and subtomogram averaging revealed an additional density near PilM in wild-type flagellar motors (Fig. 5A) that was absent in Δ*fapA* cells (Fig. 5D). Combined with the PilM-FapA interaction, this result suggests that the density may correspond to FapA. Although the overall motor architecture was resolved, including the C-ring and the periplasmic cage components PilN and PilO, the cytoplasmic component of the cage PilM and the stator region were comparatively less well resolved. These observations raise the possibility that FapA interacts with and stabilizes PilM and, directly or indirectly, influences complex stator complex dynamics. In this model, FapA and the cage could modulate stator function, thereby linking flagellar motor to changes in cellular behavior. Whether these interactions affect the motor rotational speed or torque generation and consequently contribute to the observed soft-agar migration phenotypes remains to be investigated.

*H. pylori* FapA appears to differ functionally from the best-characterized homolog, found in *V. vulnificus*. First, *H. pylori* FapA is not required for flagellar assembly, as Δ*fapA* cells retain their flagella and exhibit increased soft-agar migration relative to wild-type cells (Fig. 1 and S2). In contrast, the *V. vulnificus* Δ*fapA* mutant is aflagellate and non-migratory on soft agar (Park et al. 2016, 2019). Second, *V. vulnificus* FapA regulates motility through a pathway involving the phosphoenolpyruvate:sugar phosphotransferase system (PTS), in which glucose promotes repression of flagellar biosynthesis (Park et al. 2016). The *H. pylori* genome lacks genes encoding conventional glucose PTS components (Marais et al. 1999; Barabote and Saier 2005), and FapA-mediated repression of soft-agar migration was unaffected by supplemental glucose under the conditions tested (Fig. S4C). Third, although the predicted structure of *H. pylori* FapA contains β-strand-rich regions resembling the PF03961 FapA β-solenoid domains of *V. vulnificus* and *B. burgdorferi* FapA, the canonical PF03961 domain is not detected in *H. pylori* FapA (Fig. 6 and S4). A custom HMM generated from the *H. pylori* FapA sequence, however, expanded the list of bacteria that possess a FapA homolog, and identified numerous new candidate FapA homologs, predominantly in Campylobacterota. These findings raise the possibility that *H. pylori* FapA, and potentially other Campylobacterota homologs, form a divergent subgroup within the FapA family. Whether this divergence underlies distinct roles in flagellar biogenesis, motility, and biofilm formation across species remains to be determined.

The flagellum can function as a mechanosensory signaling platform that, in addition to driving propulsion, couples changes in mechanical load to intracellular pathways controlling cellular behavior. In high-torque complex flagellar systems such as that of *H. pylori*, the flagellar motor may be structurally adapted for mechanosensing, with accessory structures such as the PilMNO cage providing a scaffold that stabilizes the large number of stator complexes surrounding the motor. Changes in mechanical load on the rotating flagellum may be transmitted outside-in through the flagellar cage to the cytoplasm, where they modulate cellular behaviors such as motility and biofilm formation. In this model, FapA may function as a direct cytoplasmic effector within the flagellar cage mechanosensing network, linking mechanical inputs at the flagellar motor to downstream regulation of cellular behavior.

## Acknowledgements

We thank Dr. Tom Yuzvinsky of UC Santa Cruz for assistance with sample preparation and scanning electron microscopy and the U.S. Army Research Office under grant W911NF-24-1-0303 for use of the Scios 2 Dual Beam microscope. We also thank Dr. Benjamin Abrams at the UC Santa Cruz Life Sciences Microscopy Center for training and technical support with Zeiss Axio Imager (RRID: SCR_021135). This research used resources of the shared high-performance computing facility at the UCSC Hummingbird computational cluster, supported by Information Technology Services Division and the Office of Research. Cryo-ET data were collected at the Yale Cryo-EM resources funded in part by the NIH Grant 1S10OD023603-01A1. J.Y. and J.L. were supported in part by the NIAID grants R01AI198530 and R01AI087946. This research was supported in part by the NIH National Institute of Allergy and Infectious Disease (NIAID) grants RO1AI116946, RO1AI164682, and R01AI198564 to K.M.O. The funders had no role in study design, data collection and analysis, interpretation of data, or decision to submit the work for publication.

## Materials and Methods

### Bacterial strains and growth conditions

*H. pylori* G27 wild-type and derivative strains used in this study are listed in Table S1. Strains were routinely revived from frozen stocks on solid medium and cultured at 37°C under microaerobic conditions (5% O_2_, 10% CO_2_ and 85% N_2_) either in liquid medium shaking at 200 rpm or on solid medium. Solid medium consisted of Columbia Blood Agar Base (Difco) supplemented with β-cyclodextrin (0.2% w/v), cycloheximide (50 µg/ml), vancomycin (10 µg/ml), cefsulodin (5 µg/ml), polymyxin B (2.5 U/ml), and defibrinated horse blood (5% v/v) (Hemostat Labs) (CHBA). Liquid cultures were grown in Brucella broth (Difco) supplemented with 10% (v/v) heat-inactivated fetal bovine serum (Life Technologies) (BB10).

Natural transformation in *H. pylori* was performed with either PCR-amplified linear DNA, circular plasmids or genomic DNA as previously described (Rader et al. 2011; Lowenthal et al. 2009). Briefly, cells were harvested from CHBA plates and resuspended in 500 µl of BB10 medium. DNA was added to a 100-µl aliquot of cell suspension and incubated at room temperature for 5-10 min. The mixture was then spotted onto a fresh CHBA plate and incubated under microaerophilic conditions at 37°C for 16-20 h. Cells were harvested, resuspended in 500 µl of BB10 and plated onto selective CHBA agar plates containing appropriate antibiotics. Transformants were streak-purified, patched onto fresh selection CHBA plates, and stored at -80°C. Antibiotics used for selecting *H. pylori* transformants were kanamycin (15 µg/ml), chloramphenicol (13 µg/ml), and erythromycin (25 µg/ml).

*E. coli* strains used for routine cloning and other assays are listed in Table S2 and were cultured in LB medium supplemented with ampicillin (100 µg/ml), kanamycin (30 µg/ml) or chloramphenicol (20 µg/ml), unless otherwise specified.

### Recombinant DNA techniques

Standard molecular techniques were used for DNA manipulation according to the manufacturers’ instructions. DNA constructs were generated by polymerase chain reactions (PCR) using PrimeSTAR Max Ver.2 (Takara) or Q5 high-fidelity (NEB) DNA polymerases with primers synthesized by Integrated DNA Technologies (IDT), or by gene synthesis (Twist Bioscience). Cloning was performed with NEBuilder HiFi DNA Assembly Master Mix (NEB), and all constructs were verified by DNA sequencing (Genewiz). Plasmids and primer sequences used in this study are available upon request.

### Soft-agar migration assays

Soft-agar migration assays were performed using *H. pylori* strains grown on CHBA as inocula. Cells were harvested using a sterile pipette tip and inoculated into the center of soft-agar plates. Soft-agar plates were incubated at 37°C under microaerophilic conditions and migration halos were imaged using a ChemStudio PLUS imaging system (Analytik Jena), quantified with Fiji (ImageJ version 1.54p) (Schindelin et al. 2012), and the images processed using Adobe Photoshop (version 27.11.0). Soft-agar medium consisted of Brucella broth supplemented with 2.5% (v/v) heat-inactivated fetal bovine serum, cycloheximide (50 µg/ml), vancomycin (10 µg/ml), cefsulodin (5 µg/ml), polymyxin B (2.5 U/ml), trimethoprim (5 µg/ml), amphotericin B (8 µg/ml), and 0.35% (w/v) agar. Assays were performed with at least three biological replicates.

### Soft-agar pad assays

Overnight cultures of *H. pylori* were normalized to an OD_600_ of 0.10, and 5 µl of each culture was inoculated into the center of the agar pad. Agar pads were prepared by pouring molten soft-agar medium into petri dishes (100 x 15 mm) containing standard microscope slides (3 inch x 1 inch x 1 mm). Soft-agar medium used for the soft-agar pad assays have the same composition as that used in soft-agar migration assays. Images were acquired using a Zeiss AxioImager Z2 widefield microscope (Zeiss). To capture the entire culture spot, 4 x 4 tiled images were collected at 2.5x magnification with automated stitching and shading correction.

Microcolony density was quantified with Fiji (ImageJ version 1.54p) (Schindelin et al. 2012), using a modified version of a previously described method (McCall et al. 2018). Regions corresponding to the peripheral edge of the culture spot and areas beyond the colony edge were excluded to improve thresholding. Images were thresholded using the ImageJ Auto Threshold function (Kapur et al. 1985). Pixel density was measured within a 700 x 700-pixel square region and normalized to the area analyzed to calculate microcolony density (inverted gray values/mm²). Microcolony images were processed using Adobe Photoshop (version 27.11.0). Assays were performed with at least three biological replicates.

### Biofilm assays

Biofilm assays were performed in non-treated flat-bottom 96-well polystyrene microtiter plates (Costar no. 3370). Wells were inoculated with 200 µl of overnight cultures normalized to an OD_600_ of 0.15 and incubated at 37°C under microaerobic conditions without agitation. Following incubation, planktonic cells were removed and wells were washed twice with 300 µl of phosphate-buffered saline (PBS). Biofilms were stained with 200 µl of 0.1% (v/v) crystal violet for 10 min at room temperature and washed twice with 300 µl of PBS. Crystal violet bound to adherent cells was solubilized in 200 µl of 95% (v/v) ethanol, and absorbance was measured at 595 nm using a microplate reader. Assays were performed with at least three biological replicates, each consisting of at least five technical replicates.

### Bacterial adenylate cyclase two-hybrid (BACTH) assays

*E. coli* BTH101 strains harboring the plasmids used for the BACTH assay (Battesti and Bouveret 2012) are listed in Table S2. Protein-protein interactions were assessed using the BACTH system kit (Euromedex) according to the manufacturer’s instructions. Briefly, overnight cultures were diluted (1:100) into 3 ml LB medium supplemented with ampicillin (100 µg/ml), kanamycin (50 µg/ml), and IPTG (0.5 mM), and grown at 30°C with shaking (200 rpm) to an OD_600_ of 0.3-0.7. Cultures (100 µl) were mixed with 900 µl of ice-cold Z-buffer (0.06 M Na_2_HPO_4_.7H_2_O, 0.04 M NaH_2_PO_4_.H2O, 0.01 M KCl, 0.001 M MgSO_4_.7H_2_O, pH 7.0) containing freshly added β-mercaptoethanol. Cells were permeabilized with sodium dodecyl sulfate (SDS) and chloroform, and the samples were equilibrated to 30°C. Reaction was initiated by the addition of o-nitrophenyl-β-D-galactopyranoside (ONPG) and terminated with Na_2_CO_3_ after sufficient color development, and the reaction time was recorded. β-galactosidase activity was determined and reported as Miller units.

A strain carrying both pKT25-zip and pUT18C-zip, encoding leucine zipper fusions to the T25 and T18 adenylate cyclase fragments, respectively, served as the positive control, whereas a strain harboring the empty vectors pKNT25 and pUT18 served as the negative control. Assays were performed with at least four biological replicates, each consisting of three technical replicates.

### Fluorescence microscopy

Subcellular localization of PilO and FapA was determined using *H. pylori* strains carrying chromosomally integrated constructs that constitutively express mNeonGreen translational fusions of each protein (Table S1). Constitutive expression was achieved by placing the *mNeonGreen-pilO* construct, which generates an N-terminal mNeonGreen fusion to PilO (mNG-PilO), downstream of the chloramphenicol acetyltransferase (*cat*) gene, resulting in co-transcription of *cat* and *mNeonGreen*-*pilO* from the *cat* promoter. The *cat*-*mNeonGreen*-*pilO* cassette was integrated into the native *pilO* locus, replacing the endogenous *pilO* coding sequence without disrupting adjacent genes.

The *cat*-*fapA-mNeonGreen* construct generates a C-terminal mNeonGreen fusion to FapA and results in co-transcription of *cat* and *fapA-mNeonGreen* from the *cat* promoter. To generate the kanamycin-resistant construct while maintaining comparable expression, the *cat* open reading frame (ORF) was replaced with the *aphA3* ORF while retaining the same promoter. The resulting *aphA3*-*fapA-mNeonGreen* construct also generates a C-terminal mNeonGreen fusion to FapA and results in the co-transcription of *aphA3* and *fapA-mNeonGreen*. As a negative control, a *cat-mNeonGreen* cassette was generated, resulting in co-transcription of *cat* and *mNeonGreen*. Both *fapA-mNeonGreen* cassettes (FapA-mNG) and the *cat-mNeonGreen* control construct (mNG) were integrated into the McGee locus (Langford et al. 2006), the intergenic region (igR) between HPG27_0186 and HPG27_0187 (HP0203 and HP0204 in 26695 strain).

Strains were grown overnight (16-20 h) in 3 ml of BB10 medium at 37°C under microaerophilic conditions with shaking at 200 rpm. Cells from the overnight cultures were imaged using a Zeiss AxioImager Z2 widefield microscope (Zeiss) equipped with a 63x objective and processed using Zeiss ZEN lite (version 3.13) and Adobe Photoshop (version 27.11.0). Localization experiments were performed with at least three biological replicates.

### Cryo-ET specimen preparation and data acquisition

*H. pylori* cells were harvested by centrifugation at 5,000 xg for 5 min, washed with 1 ml PBS, and resuspended in approximately 50 µl PBS. A 5-µl aliquot of the suspension (OD_600_ = 0.2), with 10-nm colloidal gold fiducials, was applied to glow-discharged lacey carbon (Ted Pella) grids. Grids were blotted for 6–8 s and vitrified in liquid ethane using a GP2 plunge freezer (Leica).

Specimens were examined at −170°C on a 200-kV Titan Glacios electron microscope (Thermo Fisher Scientific). Images were acquired at a nominal magnification of 8,000x, corresponding to a pixel size of 5 Å. Tilt series spanning −48° to +48° in 3° increments were collected using SerialEM (Mastronarde 2005) and FastTomo (Xu and Xu 2021), with a nominal defocus of −4.8 µm and a cumulative dose of approximately 70 e⁻/Å^2^.

### Tomographic reconstruction and subtomogram averaging

Movie frames were motion-corrected using MotionCor2 (Zheng et al. 2017) and assembled into tilt stacks in IMOD (Mastronarde and Held 2017). Tilt series were aligned using gold fiducials or fiducial-free alignment. Defocus values were estimated with Gctf (Zhang 2016), followed by contrast transfer function correction using IMOD’s ctfphaseflip. Tomograms were reconstructed in Tomo3D (Agulleiro and Fernandez 2015) using simultaneous iterative reconstruction (SIRT) for motor identification and weighted back-projection (WBP) for subtomogram averaging.

A total of 192 flagellar motor particles were manually identified from 96 tomograms. Their positions and approximate orientations were used for initial alignment in bin4 tomograms. Subtomograms were then extracted from WBP reconstructions using the refined coordinates. Iterative alignment and multi-reference classification were performed in Protomo as described previously (Liu, Tachiyama, et al. 2024; Schoichi Tachiyama et al. 2022) using bin4 subtomograms, followed by further refinement with bin2 data.

### Scanning electron microscopy (SEM)

Overnight liquid cultures were normalized to an OD_600_ of 0.15 and 3 ml of each culture was inoculated into 6-well plates containing 12-mm round glass coverslips (Costar) positioned against the side of each well. Plates were incubated at 37°C under microaerophilic conditions for 20 h. Coverslips were gently removed, washed twice with PBS, and cells were fixed in 2.5% (v/v) glutaraldehyde diluted in 0.1 M HEPES for 1 h at 4°C. Samples were subsequently dehydrated through a graded ethanol series (25% v/v, 50%, 75%, 90%, 100%). Dehydrated samples were critical point dried (Balzers Union 342/11 120B) to preserve sample morphology and sputter coated with ∼20 nm of gold using a Hummer IV sputter coater (Technics Inc.). Samples were imaged using a Scios 2 Dualbeam FIB-SEM microscope (Thermo Fisher Scientific). SEM were performed with at least three biological replicates.

### Phylogenetic tree construction and homology search

The genome set used for phylogenomic analyses comprised the representative flagellated bacterial genomes reported by Feng et al. (Feng et al. 2026). Phylogenetic trees of representative flagellated bacteria and flagellated Campylobacterota were constructed using GToTree (Lee 2019) with the prepackaged Bacteria and Epsilonproteobacteria conserved single-copy genes hidden Markov models (HMMs). Of the original 1,365 representative flagellated genomes, *Candidatus Chryseopegocella kryptomonas* (GCF_001536065.1) did not meet the minimum threshold for conserved bacteria HMMs hits and was excluded from the analysis. Concatenated multiple sequence alignments were used to construct maximum-likelihood phylogenetic trees using IQ-TREE 3 (Wong et al. 2026). The best fit model (LG + R10) was selected by ModelFinder (Kalyaanamoorthy et al. 2017), and branch support was assessed with 1,000 bootstrap replicates. Trees were visualized and annotated using iTOL v6 (Letunic and Bork 2024).

To identify FapA homologs across the representative flagellated bacterial genomes, we used a Pfam profile HMM corresponding to the FapA β-solenoid domain (PF03961) and a custom HMM corresponding to the putative Campylobacterota-specific FapA β-solenoid domain. The profile HMM for the putative Campylobacterota-specific FapA β-solenoid domain was constructed from homologs of the *H. pylori* G27 FapA (HPG27_837) residues 300-611 identified by PSI-BLAST (Altschul 1997). Homologous sequences were aligned using MAFFT (Katoh 2002) and the resulting alignment was trimmed with trimAl (Capella-Gutiérrez et al. 2009) before construction of the profile HMM using hmmbuild (Johnson et al. 2010).

Proteomes of the representative flagellated bacterial and Campylobacterota proteomes were retrieved from NCBI and concatenated into a unified proteome. Profile HMM searches were performed against this database with hmmsearch (E-value < 1e^-3^) (Johnson et al. 2010).

### Protein structure prediction

Protein structural predictions were carried out using AlphaFold 3 (Abramson et al. 2024) and visualized using UCSF ChimeraX (version 1.10.1) (Pettersen et al. 2021).

## Statistical analysis

Statistical analysis was performed using one-way analysis of variance (ANOVA) followed by Tukey’s multiple-comparison test or Student’s *t*-test. All analyses were carried out using GraphPad Prism software (version 11).

