## Supplemental for "FapA extends the flagellar cage-dependent network into the cytoplasm to control surface-associated behaviors in *Helicobacter pylori*"

The PDF file includes:

Tables S1, S2

Figures S1-S4

**Table S1.** *H. pylori* strains used in this study.

| Strain | Strain # | Relevant genotype/description | Antibiotic resistance[a] | Reference /source |
|---|---|---|---|---|
| **Deletion strains** | | | | |
| WT | KO379 | G27 wild type | None | (Censini et al. 1996) |
| Δ*fapA* | KO2037 | Δ*fapA::aphA3* | Kan | This study |
| Δ*fapA::cat* | KO2136 | Δ*fapA::cat* | Cm | This study |
| Δ*pilM* | KO1741 | Δ*pilM::aphA3* | Kan | (Liu et al. 2024) |
| Δ*fapA* Δ*pilM* | KO2158 | Δ*fapA::cat* Δ*pilM::aphA3* | Cm, Kan | This study |
| Δ*pilMNO* | KO1743 | Δ*pilMNO::aphA3* | Kan | (Liu et al. 2024) |
| Δ*fapA* Δ*pilMNO* | KO2137 | Δ*fapA::cat* Δ*pilMNO::aphA3* | Cm, Kan | This study |
| Δ*motAB* | KO1538 | Δ*motAB* | None | This study |
| Δ*pilMNO* Δ*motAB* | KO2204 | Δ*pilMNO::aphA3* Δ*motAB* | Kan | This study |
| Δ*fliG* | KO1063 | Δ*fliG::cat* | Cm | (Lertsethtakarn et al. 2015) |
| Δ*pilMNO* Δ*fliG* | KO2393 | Δ*pilMNO::erm* Δ*fliG::cat* | Cm, Erm | This study |
| Δ*motAB* Δ*fliG* | KO2261 | Δ*motAB* Δ*fliG::cat* | Cm | This study |
| Δ*pilMNO* Δ*motAB* Δ*fliG* | KO2392 | Δ*pilMNO::erm* Δ*motAB* Δ*fliG::cat* | Cm, Erm | This study |
| Δ*fliL* | KO2259 | Δ*fliL::cat* | Cm | This study |
| Δ*cheA* | KO857 | Δ*cheA::cat* | Cm | (Lertsethtakarn et al. 2015) |
| Δ*fapA* Δ*cheA* | KO2199 | Δ*fapA::aphA3* Δ*cheA::cat* | Cm, Kan | This study |
| Δ*cheV1* | KO1277 | Δ*cheV1::cat* | Cm | (Lertsethtakarn and Ottemann 2010) |
| Δ*fapA* Δ*cheV1* | KO2260 | Δ*fapA::aphA3* Δ*cheV1::cat* | Cm, Kan | This study |
| Δ*cheZ* | KO1315 | Δ*cheZ* | None | (Lertsethtakarn et al. 2015) |
| Δ*fapA* Δ*cheZ* | KO2200 | Δ*fapA::aphA3* Δ*cheZ* | Kan | This study |
| **Localization strains** | | | | |
| WT (mNG) | KO2350 | G27 containing *cat-mNeonGreen* cassette in the intergenic region (igR) | Cm | This study |
| WT (mNG-PilO) | KO2139 | G27 containing *cat-mNeonGreen-pilO* in the native *pilO* locus | Cm | This study |
| WT (FapA-mNG) | KO2344 | G27 containing *cat-fapA-mNeonGreen* in the igR | Cm | This study |
| WT (Kan-FapA-mNG) | KO2348 | G27 containing *aphA3-fapA-mNeonGreen* in the igR | Kan | This study |
| Δ*fapA* (mNG) | KO2351 | Δ*fapA::aphA3* containing *cat-mNeonGreen* cassette in the igR | Cm, Kan | This study |
| Δ*fapA* (FapA-mNG) | KO2297 | Δ*fapA::aphA3* containing *cat-fapA-mNeonGreen* cassette in the igR | Cm, Kan | This study |

**Table S1.** *H. pylori* strains used in this study (continue).

| Strain | Strain # | Relevant genotype/description | Antibiotic resistance[a] | Reference /source |
|---|---|---|---|---|
| **Localization strains** (continue) | | | | |
| Δ*fapA* (Kan-FapA-mNG) | KO2456 | Δ*fapA::cat* containing *aphA3-fapA-mNeonGreen* cassette in the igR | Cm, Kan | This study |
| Δ*pilMNO* (FapA-mNG) | KO2299 | Δ*pilMNO::aphA3* containing *cat-fapA-mNeonGreen* cassette in the igR | Cm, Kan | This study |
| Δ*motAB* (FapA-mNG) | KO2390 | Δ*motAB* containing *cat-fapA-mNeonGreen* cassette in the igR | Cm | This study |
| Δ*fliG* (Kan-FapA-mNG) | KO2349 | Δ*fliG::cat* containing *aphA3-fapA-mNeonGreen* cassette in the igR | Cm, Kan | This study |
| Δ*pilMNO* Δ*motAB* (FapA-mNG) | KO2450 | Δ*pilMNO::aphA3* Δ*motAB* containing *cat-fapA-mNeonGreen* cassette in the igR | Cm, Kan | This study |
| Δ*motAB* Δ*fliG* (kan-FapA-mNG) | KO2354 | Δ*motAB* Δ*fliG::cat* containing *aphA3-fapA-mNeonGreen* cassette in the igR | Cm, Kan | This study |
| Δ*pilMNO* Δ*motAB* Δ*fliG* (Kan-FapA-mNG) | KO2451 | Δ*pilMNO::erm* Δ*motAB* Δ*fliG::cat* containing *aphA3-fapA-mNeonGreen* cassette in the igR | Cm, Erm, Kan | This study |
| Δ*fliF* (Kan-FapA-mNG) | KO2455 | Δ*fliF::cat* containing *aphA3-fapA-mNeonGreen* cassette in the igR | Cm, Kan | This study |
| Δ*cheA* (Kan-FapA-mNG) | KO2353 | Δ*cheA::cat* containing *aphA3-fapA-mNeonGreen* cassette in the igR | Cm, Kan | This study |
| Δ*cheZ* (FapA-mNG) | KO2345 | Δ*cheZ* containing *cat-fapA-mNeonGreen* cassette in the igR | Cm | This study |

[a]Kan, kanamycin; Cm, chloramphenicol; Erm, erythromycin.

**Table S2.** *E. coli* strains and plasmids used in this study.

| Strain/plasmid | Strain # | Relevant genotype/description | Antibiotic resistance[a] | Reference/ source |
|---|---|---|---|---|
| **Strains** | | | | |
| DH5α | KO133 | *endA1*, *hsdR17*, (rK$^-$, mK$^+$), *supE44*, *thi*$^-$1, *recA1*, *gyrA*, Nal$^R$, *relA1*, Δ(*lacZYA-argF*), U169, *deoR*, fφ80Δ(*lac*Z)M15 | Nal | NEB |
| BTH101 | KO1350 | F$^-$, *cya-99, araD139, galE15, galK16, rpsL1* (Str$^R$), *hsdR2, mcrA1, mcrB1* | Str | Euromedex |
| **Plasmids** | | | | |
| pUC19 | KO137 | Cloning plasmid | Amp | (Yanisch-Perron et al. 1985) |
| pΔ*fapA::cat* | KO2132 | pUC19 containing Δ*fapA::cat* | Amp, Cm | This study |
| **Plasmids for BACTH assays** | | | | |
| pKNT25 | KO1345 | BACTH cloning plasmid | Kan | Euromedex |
| pKT25 | KO1344 | BACTH cloning plasmid | Kan | Euromedex |
| pUT18 | KO1346 | BACTH cloning plasmid | Amp | Euromedex |
| pUT18C | KO1347 | BACTH cloning plasmid | Amp | Euromedex |
| pKT25-zip | KO1348 | BACTH positive control plasmid with leucine zipper | Kan | Euromedex |
| pUT18C-zip | KO1349 | BACTH positive control plasmid with leucine zipper | Amp | Euromedex |
| pKNT25-*pilM* | KO2065 | pKNT25 containing *pilM*-T25 translational fusion | Kan | This study |
| pKT25-*pilM* | KO2068 | pKT25 containing T25-*pilM* translational fusion | Kan | This study |
| pUT18-*pilM* | KO2074 | pUT18 containing *pilM*-T18 translational fusion | Amp | This study |
| pUT18C-*pilM* | KO2071 | pUT18C containing T18-*pilM* translational fusion | Amp | This study |
| pKNT25-*fapA* | KO2066 | pKNT25 containing *fapA*-T25 translational fusion | Kan | This study |
| pKT25-*fapA* | KO2069 | pKT25 containing T25-*fapA* translational fusion | Kan | This study |
| pUT18-*fapA* | KO2075 | pUT18 containing *fapA*-T18 translational fusion | Amp | This study |
| pUT18C-*fapA* | KO2072 | pUT18C containing T18-*fapA* translational fusion | Amp | This study |
| **Plasmids for protein localization** | | | | |
| pG27-igR::*cat-mNeonGreen* | KO2332 | pUC19 derivative containing *cat-mNeonGreen* cassette flanked by intergenic region (igR) fragments (mNG) | Amp, Cm | This study |
| pG27-igR::*cat-fapA-mNeonGreen* | KO2289 | pUC19 derivative containing *cat-fapA-mNeonGreen* cassette flanked by igR fragments (FapA-mNG) | Amp, Cm | This study |
| pG27-igR::*aphA3-fapA-mNeonGreen* | KO2331 | pUC19 derivative containing *aphA3-fapA-mNeonGreen* cassette flanked by igR fragments (Kan-FapA-mNG) | Amp, Kan | This study |
| pG27-*pilO*::*cat-mNeonGreen-pilO* | KO2133 | pUC19 derivative containing *cat-mNeonGreen-pilO* cassette flanked by *pilO* up- and downstream fragments (mNG-PilO) | Amp, Cm | This study |

[a]Nal, nalidixic acid; Str, streptomycin; Amp, ampicillin; Kan, kanamycin; Cm, chloramphenicol.

### Supplementary Figures

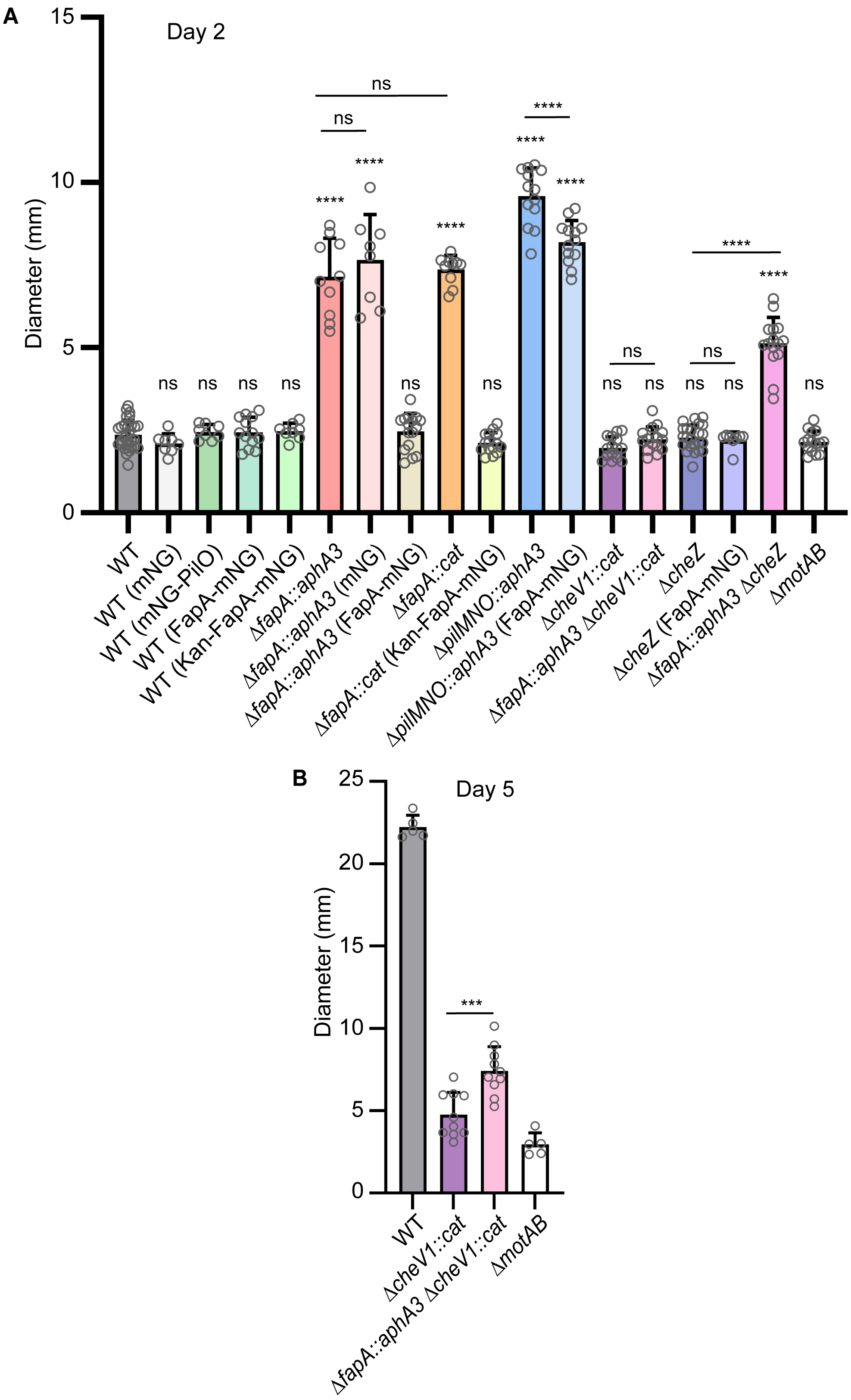

**C**

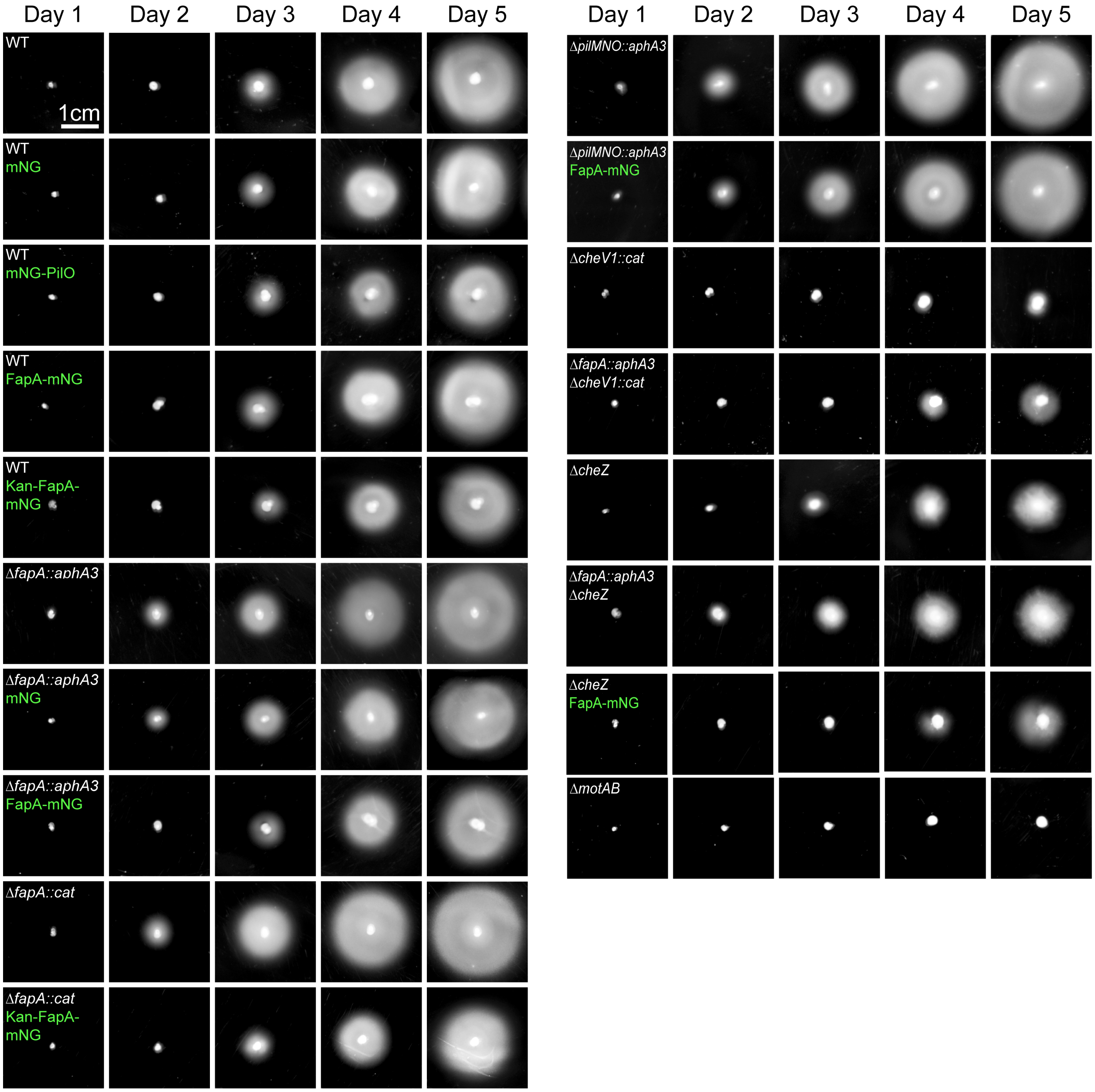


**Fig. S1. FapA-mNeonGreen fusions complement the enhanced soft-agar migration phenotype of Δ*fapA*.** (A) Migration zone diameters measured on (A) day 2 and (B) day 5, and (C) representative images of migration zones on day 1-5 for G27 wild type and the indicated mutants with or without chromosomal copies of unfused mNeonGreen (mNG), FapA-mNG, or mNG-PilO translational fusions. The repression of soft-agar migration exhibited by G27 wild type on day 2 was not affected by the presence of fluorescent constructs. The Δ*fapA::aphA3* with or without the mNG control construct exhibited enhanced migration on day 2. Δ*fapA::cat* strain exhibited similar enhanced migration as Δ*fapA::aphA3* strain. Expression of FapA-mNG and Kan-FapA-mNG abolished the enhanced migration of Δ*fapA::aphA3* and Δ*fapA::cat*, respectively, restoring migration to a level similar to that of the wild type, indicating that the FapA-mNG fusion proteins are functional. The cage mutant Δ*pilMNO::aphA3* expressing FapA-mNG exhibited reduced migration compared to the parental Δ*pilMNO::aphA3* strain, consistent with FapA-mediated repression of migration. Deletion of Δ*fapA* increased migration in the CW-biased Δ*cheZ* background on day 2 and the CCW-biased Δ*cheV1* background on day 5, indicating that FapA-mediated repression of soft-agar migration is independent of flagellar rotational bias. Δ*motAB* served as a nonmotile control. mNG, FapA-mNG and Kan-FapA-mNG fluorescent constructs were integrated into the McGee locus, the intergenic region (igR) between HPG27_0186 and HPG27_0187 (HP0203 and HP0204 in 26695 strain), whereas the mNG-PilO construct was integrated at the native *pilO* locus. Each data point represents a technical replicate from at least two independent biological replicates. Statistical significance was determined using one-way ANOVA followed by Tukey's multiple-comparisons test for day 2 and an unpaired Student's *t*-test for day 5. Error bars represent standard deviations. ****, $p < 0.0001$; ***, $p = 0.0005$; ns, not significant. CW, clockwise; CCW, counterclockwise; mNG, mNeonGreen with a *cat* selectable marker; mNG-PilO, mNeonGreen-PilO translational fusion with a *cat* selectable marker; FapA-mNG, FapA-mNeonGreen translational fusion with a *cat* selectable marker; Kan-FapA-mNG, FapA-mNeonGreen translational fusion with an *aphA3* selectable marker.

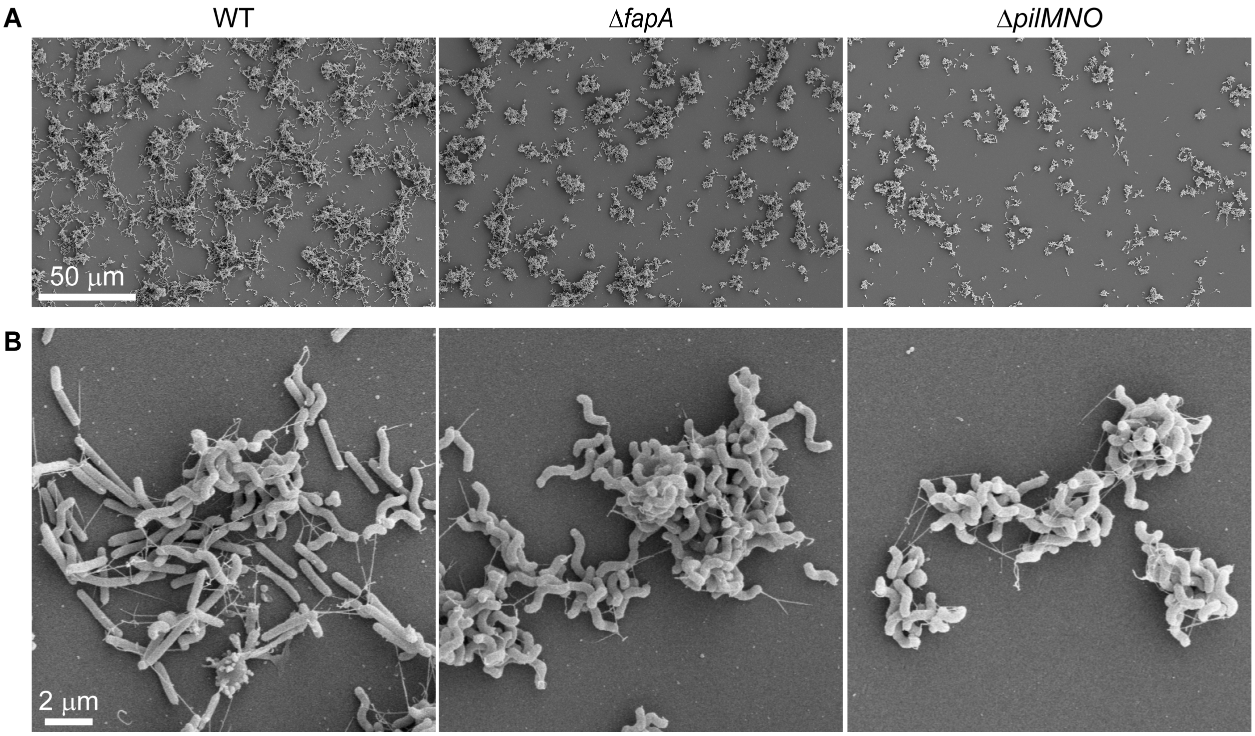


**Fig. S2. Δ*fapA* retains flagella and exhibits biofilm and cell aggregation phenotypes intermediate between those of G27 wild type and the cage mutant.** Scanning electron microscopy (SEM) images of G27 wild type, Δ*fapA*, and Δ*pilMNO* after 20 hours of growth, showing (A) distinct biofilm and cell aggregation architectures and (B) higher-magnification views of cell aggregates and flagellar filaments. Both Δ*fapA* and Δ*pilMNO* retained their flagella, indicating that their altered biofilm and cell aggregation phenotypes are not due to loss of flagellation. Images are representative of at least three biological replicates.

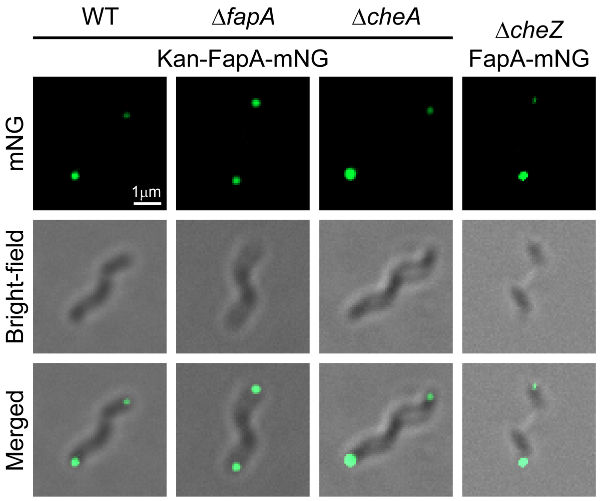


**Fig. S3. FapA-mNeonGreen polar localization is retained with an alternative construct carrying the *aphA3* selectable marker.** FapA-mNeonGreen expressed from a construct carrying the *aphA3* selectable marker (Kan-Fap-mNG) localizes to the poles in G27 wild-type and *ΔfapA* strains, similar to FapA-mNeonGreen expressed from a construct carrying the *cat* selectable marker (FapA-mNG). FapA also localizes to the cell poles in Δ*cheA* (CCW-biased) and Δ*cheZ* (CW-biased) strains, suggesting that changes in flagellar rotation bias do not disrupt FapA polar localization. Images shown are representative of at least three biological replicates.

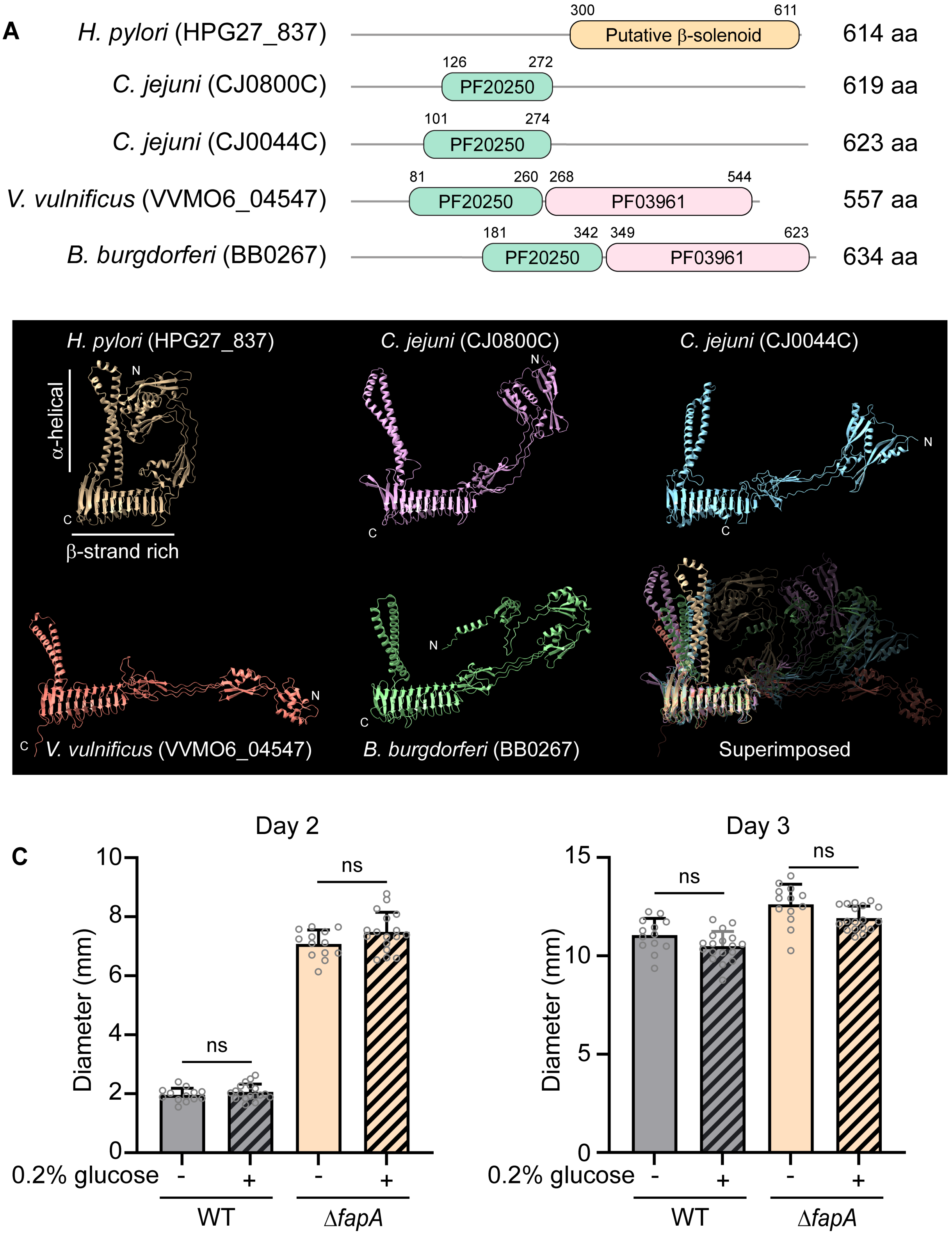


**Fig. S4. Comparison of domain architectures and predicted structures of FapA homologs.** (A) Domain architectures and (B) predicted structures of FapA homologs from *H. pylori*, *C. jejuni*, *V. vulnificus* and *B. burgdorferi*. Domain annotations for the proteins from *C. jejuni* CJ0800C (UniProt: Q0PA90) and CJ0044C (UniProt: Q0PC82), *V. vulnificus* VVMO6_04547 (UniProt: A0A455ZM43), and *B. burgdorferi* BB0267 (UniProt: H7C7R5) were retrieved from UniProt. PF03961, FapA β-solenoid domain; PF20250, FapA N-terminal region. Residues 300-611 of *H. pylori* FapA (HPG27_837; UniProt: B5Z7P3), encompassing predicted β-strand-rich and α-helical regions (indicated in the *H. pylori* model), were used to generate a custom hidden Markov model (HMM) to identify candidate FapA homologs containing a putative Campylobacterota-specific β-solenoid region. N- and C-termini of the predicted protein structures are indicated. The final panel shows structural superimposition of the five predicted models, aligned based on their β-solenoid regions. Structural predictions were generated using AlphaFold3 and visualized using UCSF ChimeraX. (C) Soft-agar migration assays showing migration diameters of wild-type G27 and Δ*fapA* strains on days 2 and 3 in soft agar with or without supplemental 0.2% (w/v) glucose. No significant differences in migration were observed between glucose-supplemented and unsupplemented conditions for either strain.
